# Genome-wide CRISPR screening identifies new regulators of glycoprotein secretion

**DOI:** 10.1101/522334

**Authors:** Stephanie J. Popa, Julien Villeneuve, Sarah Stewart, Esther Perez Garcia, Anna Petrunkina Harrison, Kevin Moreau

## Abstract

The fundamental process of protein secretion from eukaryotic cells has been well described for many years, yet gaps in our understanding of how this process is regulated remain. With the aim of identifying novel genes involved in the secretion of glycoproteins, we used a screening pipeline consisting of a pooled genome-wide CRISPR screen followed by secondary siRNA screening of the hits to identify and validate several novel regulators of protein secretion. We present approximately 50 novel genes not previously associated with protein secretion, many of which also had an effect on the structure of the Golgi apparatus. We further studied a small selection of hits to investigate their subcellular localisation. One of these, GPR161, is a novel Golgi-resident protein that we propose maintains Golgi structure via an interaction with golgin A5.

## INTRODUCTION

Protein secretion is a fundamental and well-known process in cell biology, in which proteins are transported from the endoplasmic reticulum (ER) to the Golgi via COP coated vesicles, and subsequently to the plasma membrane^1–3^. The localisation and activity of proteins involved in this secretory pathway must be tightly regulated to ensure correct spatio-temporal distribution of membranes and cargo proteins along the pathway. While the molecular machinery of secretion is relatively well understood, our knowledge remains incomplete, particularly regarding the regulation of protein secretion. For example, evidence suggests that multiple trafficking routes for transport within the Golgi exist^4^. The complex process of sorting proteins at the Golgi into vesicles for their correct destinations also requires further investigation^5–7^.

Recently, genome-wide CRISPR screening has emerged as a powerful strategy to identify novel gene functions; this type of screening has the advantages of being unbiased and more reliable than previously-used methods, in terms of introducing genetic mutations^8^. As such, it provides an opportunity to uncover new information about the regulation of protein secretion. Pooled CRISPR screening has been used for a wide range of applications, including the investigation of drug-resistance mechanisms in cancer cells^9^, the genetics of pluripotency^10^, autophagy regulators^11,12^ and host factors required for viral infection^13^. We previously demonstrated that unbiased pooled genome-wide CRISPR screening could effectively reveal key players required for glycoprotein secretion, using galectin-3 retention at the cell surface to assess glycoprotein secretion^14^. Galectin-3 is a cytosolic protein that is secreted without entering the conventional secretory pathway for its export to the extracellular space. Vesicular and non-vesicular modes of secretion have been proposed, but the precise series of events involved in the unconventional secretion of galectin-3 remain ill defined^15^. Once outside the cell, galectin-3 binds to β-galactosides resident on the cell surface^16^ and can therefore be used as an indirect measure of glycoprotein transport to the cell surface via the ER-Golgi secretory pathway. Using a binding partner of glycans as a readout, rather than one specific glycoprotein, allows regulators of general glycoprotein secretion to be discovered.

Here we devised a powerful experimental pipeline, using an improved pooled genome-wide CRISPR screen followed by two arrayed secondary screening methods using siRNA knockdown to identify new factors involved in glycoprotein secretion and Golgi apparatus architecture. Combining these methods allows the speed and reduced cost of pooled CRISPR screening to be taken together with the advantages of an arrayed siRNA screen in which genotype and phenotype remain linked^17^. Using this approach, we were able to validate 55 novel hits that are important for glycoprotein secretion. We found that many of these hits are also important for maintenance of the Golgi architecture. One of these hits, GPR161, is a novel Golgi-localised protein that decreases glycoprotein secretion and disrupts the Golgi structure.

## RESULTS

### A genome-wide CRISPR screen revealed novel regulators of glycoprotein secretion

We carried out a genome-wide CRISPR screen with the aim of identifying novel genes involved in glycoprotein secretion, using the level of galectin-3 on the surface of live cells to look at glycoprotein secretion via the ER-Golgi pathway. The workflow for the screen is shown in figure 1A. Suspension HeLa cells stably expressing Cas9 were transduced with the Brunello lentiviral guide (sgRNA) library, an optimised library for the human genome^18^, at a low multiplicity of infection such that the majority of the transduced cells received exactly one sgRNA. After transduction, we performed three rounds of cell sorting by fluorescence activated with increasing selection stringency for low levels of cell surface galectin-3. This method initially allows more cells through cell sorting, then becomes gradually more stringent by decreasing the gating percentage, increasing the number of true positives in later sorted populations. This resulted in the enrichment of two distinct populations with low levels of cell-surface galectin-3 after the third sort, in population N3 (figure 1B). As there are lots of cells with very little or no galectin-3 on the cell surface in population N3, the other populations may have more galectin-3, transferred from the extracellular pool, or a higher effective concentration of anti-galectin-3 antibody. A fourth round of cell sorting was used to separate these distinct populations, resulting in populations M4 and N4 (Figure 1B). We extracted genomic DNA from all populations and carried out deep sequencing to determine the raw sgRNA counts in each population, available in supplementary data S1. MaGECK analysis allowed the identification of hits from population N1 onwards; full MaGECK results are available in supplementary data S2. Hits enriched in populations N1 and N4 are shown in figure 1C.

**Figure 1:**
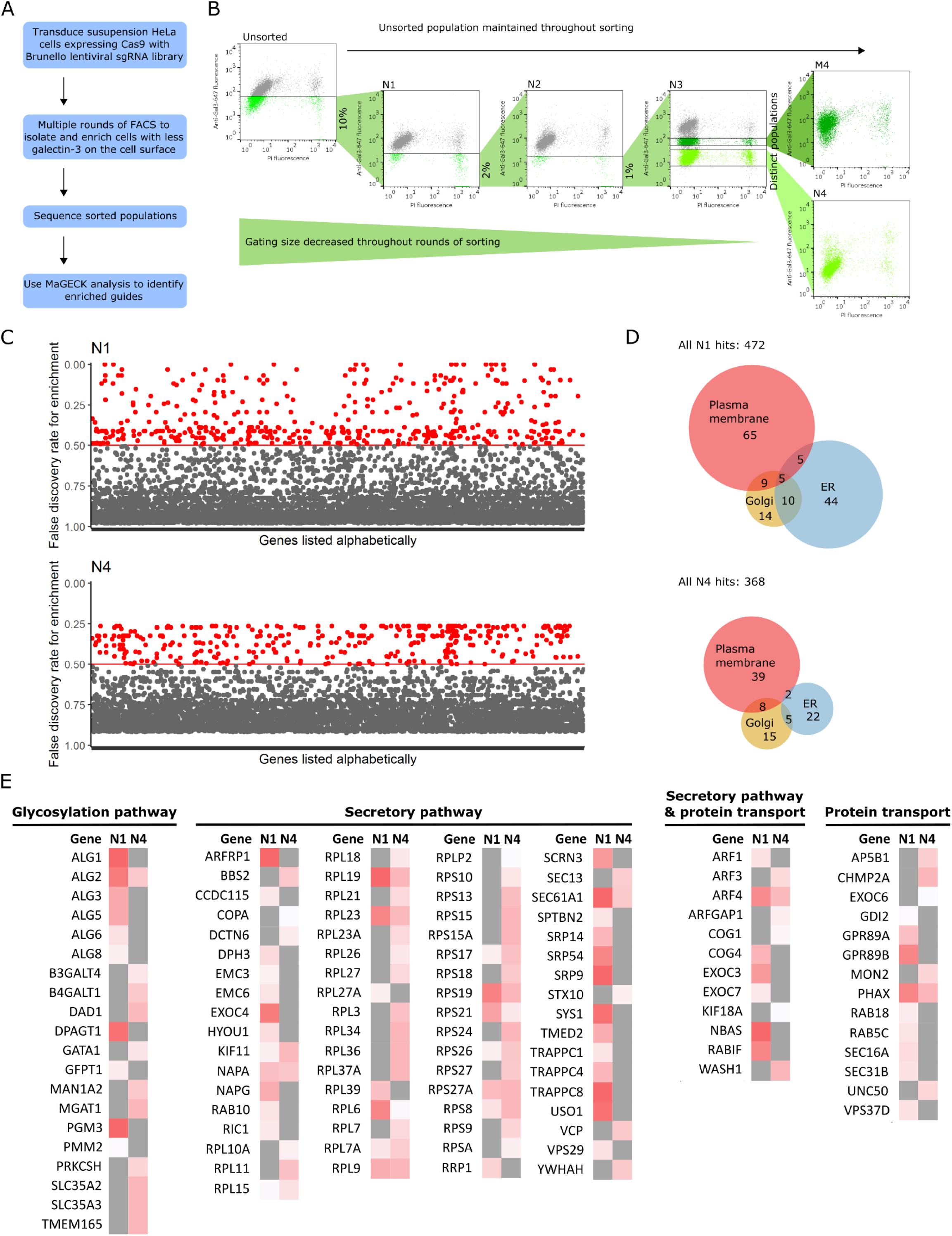
CRISPR screen. (A) Workflow of genome-wide CRISPR screen. (B) Multiple rounds of FACS sorting, decreasing gating percentage with progressive rounds, resulted in final populations of cells with a low level of galectin-3 on the cell surface. FACS plots are of samples taken after cells had been expanded after sorting, and show cell surface galectin-3 levels as measured by anti-galectin-3-647 fluorescence, and viability as measured by PI. Labels of FACS plots refer to population names: N1-4 = populations after negative sorts 1-4; M4, cells expressing medium level of galectin-3 after sort 4. (C) Hits from populations N1 and N4; genes with a false discovery rate (FDR) <0.5 were defined as hit and shown in red. Genes with had guides that were enriched, but not significantly enriched (i.e. FDR ≥ 0.5), are shown in grey. (D) Euler diagram showing the number of hits known to be associated with organelles in the secretory pathway. Many ER-resident hits are lost when progressing from sorts N1 to N4. (E) Heat map of genes known to be in the secretory pathway, glycosylation pathway or involved in protein transport, as annotated by GO_Slim lists. Grey indicates an FDR greater than 0.5; hits are shown on a white-red scale, where darker red indicates a more significant FDR, closer to 0. The identity of hits changes significantly from populations N1 to N4.

The identity of the hits changed as cell sorting progressed. This was particularly true of hits which localise to the ER: in total, 64 hits localise to the ER in population N1, compared to 29 in population N4 (figure 1D). As many genes in the ER are involved in critical biological processes, it is likely that these genes were lost in later sorts as cells lacking these genes have a growth or survival defect. For example, many of the enzymes involved in N-linked glycosylation, such as the asparagine-linked glycosylation (ALG) enzymes ALG1, ALG3, ALG5, ALG6 and ALG8, are identified as hits in population N1 but are lost in N4. Conversely, some hits are enriched after multiple cell sorting rounds in N4 but were not identified in N1 (figure 1E). As such, we took a selection hits from across all populations forward into the secondary screen. MaGECK-MLE and MaGECK-RRA analysis revealed that there is little difference between the later populations: N3, M4 and N4 (supplementary data S2). As we intended to validate hits with further screening, we used a false discovery rate (FDR) of 0.5 to classify hits. This lenient FDR is validated by the presence of hits previously shown to play a role in secretory and glycosylation pathways, such as ribosomal proteins involved in protein translocation into the ER machinery, enzymes involved in N-linked glycosylation, RAB proteins and trafficking protein particle complex (TRAPPC) proteins (figure 1E). This shows that our screen was very efficient in identifying previously described regulators of glycoprotein secretion. Additionally, there were many hits identified that did not have a clear link to functions in protein secretion. Therefore, we hypothesised that many genes enriched in our screen with either no characterised function, or with no described function in the secretory pathway, may be new regulators of protein secretion.

### An RNAi secondary screen validated hits from the CRISPR screen

Due to the lenient FDR used for MaGECK it was important to validate the hits. Around 400 hits, identified from populations N1 - N4/M4 of the CRISPR screen, were selected for validation by secondary screening. 113 of these 400 hits had no annotated function in UniProt; 32 also had no gene ontology (GO) annotations. As many of these hits were uncharacterised, they had the potential to represent novel regulators of protein secretion. Twelve positive controls annotated with GO terms related to protein secretion were also included in the screen to validate the statistical analysis. For secondary screening, we used siRNA knockdown of hits in HeLa cells that stably express horseradish peroxidase fused to a signal sequence (ss-HRP), which is a glycosylated protein secreted into the extracellular space^19^. HeLa-ss-HRP were transfected with arrayed siRNA in a 96 well plate. 72 h after transfection, we collected the supernatant, lysed the cells and assessed HRP by chemiluminescence. Knockdown that led to a decrease in the ratio of supernatant luminescence to cell lysate luminescence indicated genes that were important for glycoprotein secretion. Knockdown of 92 of the genes screened resulted in less secretion of HRP (figure 2A). Positive controls also resulted in less HRP secretion, as expected (figure 2A-B). The 92 genes validated by the siRNA screen are listed in figure 2B-C. Of these hits, 55 were not annotated with GO_Slim terms related to secretion or protein transport; 22 of these had no GO_Slim annotations and 6 had no GO annotations at all (figure 2C).

**Figure 2:**
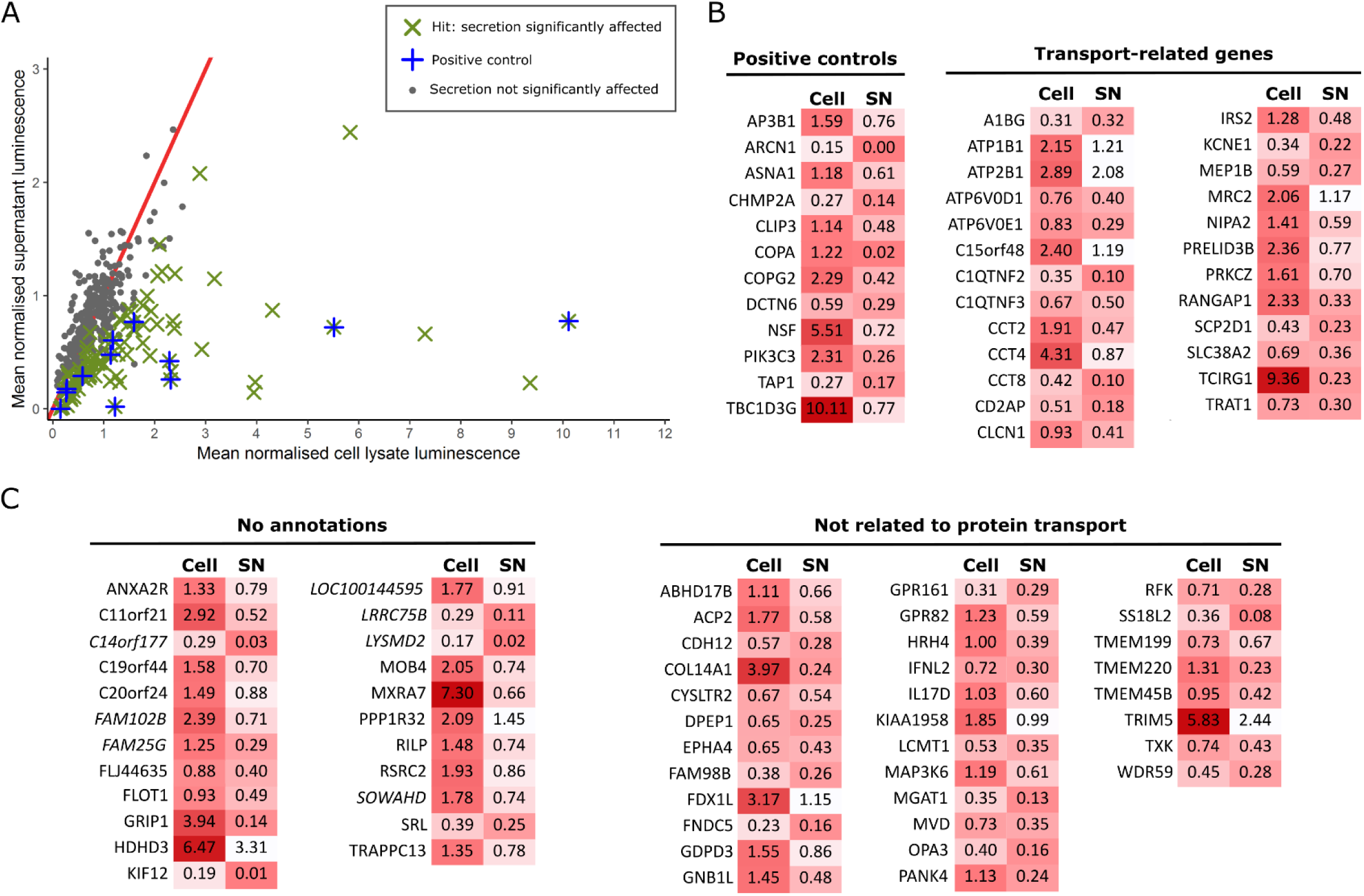
HRP screen to identify genes important for secretion. Approximately 400 genes were screened in an arrayed siRNA knockdown format. (A) Mean of the normalised luminescence for both cell lysate and supernatant signals from two independent replicates. ∼400 genes were screened; those with significant divergence from control siRNAs, as identified by the ROUT method, are indicated by a green cross (**x**). Twelve genes known to be involved in secretion or protein trafficking were included in the screen as positive controls, indicated by a blue plus (**+**); these known controls were also identified as hits, validating the statistical method used here. (B-C) Mean values for normalised luminescence for cell lysate and supernatant (SN) signals of the 92 hits identified. Here these hits are shown on a white-red scale, where darker red indicates a greater secretion defect. The generic GO_Slim was used to categorise hits as either related to protein transport, unrelated to protein transport or not annotated. (B) A gene was defined as related to protein transport if it was annotated with any of the GO_Slim annotations transport (GO:0006810), transmembrane transport (GO:0055085) or vesicle-mediated transport (GO:0016192). (C) Genes unrelated to protein transport or not annotated by GO_slim terms represent novel hits involved in protein secretion. Of the genes with no GO_Slim annotations, genes with no GO annotations at all are indicated in italics. Genes defined as not related to protein transport have other annotations in the generic GO_Slim.

### Further screening revealed many genes with altered glycoprotein secretion also had a fragmented Golgi apparatus

To further probe how the identified genes affect glycoprotein secretion, the 92 genes identified in the HRP screen were further analysed for changes in Golgi apparatus morphology using the same siRNA knockdown. 72 h after siRNA transfection, cells were fixed and analysed by immunofluorescence, using an antibody against GM130, a marker of *cis*-Golgi membranes. Images were collected and classified as having fragmented or intact *cis-*Golgi using the machine learning platform within CellProfiler as shown in figure 3A-B. Many of the silenced genes resulted in an increase in fragmented *cis-*Golgi compared to the control of ∼20%. Genes with more than the median percentage of cells with fragmented Golgi are shown in figure 3C, and all genes are shown in supplementary figure S3. Based on having unknown functions in secretion, five genes from the top half of these hits were chosen to further investigate the mechanism by which these hits lead to a secretion defect: *GPR161, TMEM220, FAM98B, FAM102B* and *MXRA7*. We confirmed that the knockdown of these five genes led to a perturbation in the architecture of the Golgi and affected protein secretion. This can be seen by the more diffuse localisation of two Golgi-resident proteins, TGN46 and GM130; by the more diffuse localisation of SEC31, a marker of ER exit sites, and by the defect in the transport of the glycoprotein MHC-I to the cell surface (figure 3D).

**Figure 3:**
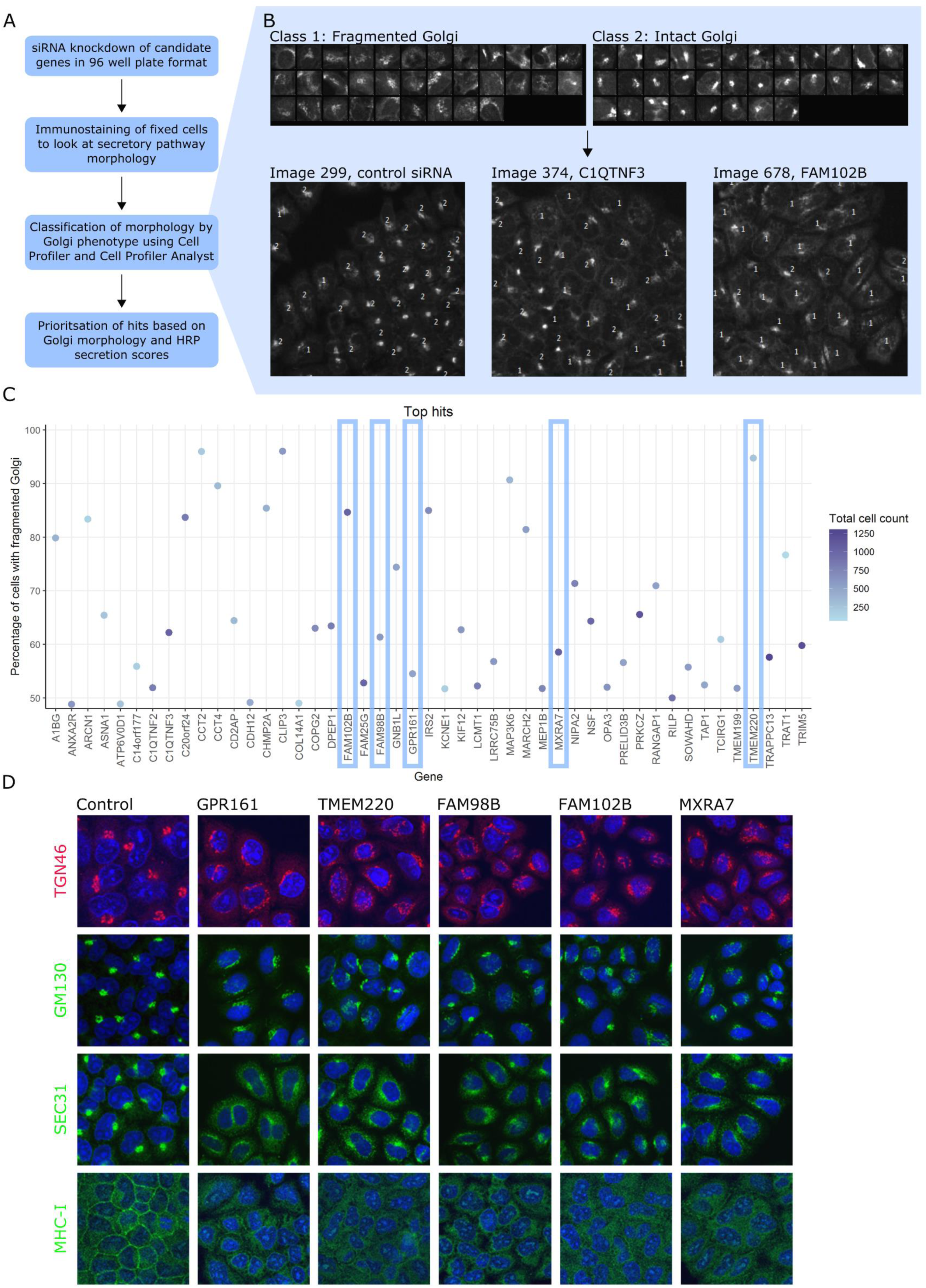
Further secondary screen for altered Golgi morphology. (A) Workflow of Golgi morphology screen. (B) Training set used to train CPA’s classifier to distinguish between fragmented and intact Golgi. Using this training set, sensitivity was 77 % for fragmented Golgi class and 82 % for the intact Golgi class. Examples of control and test images classified using CPA are also shown. Cells with fragmented Golgi are labelled with 1; cells with intact Golgi are labelled with 2. (C) Percentage of cells with fragmented Golgi for the top 48 hits from tertiary screen. Hits are arranged alphabetically and coloured by cell count, with darker blue spots representing more confluent wells. Five of these hits were further characterised, indicated with blue rectangles. (D) Immunostaining of HeLa cells treated with siRNA to knockdown each of the five genes indicated above the images. Cells were stained with the antibodies against the proteins indicated in green or red on the left. For all images, DAPI staining is shown in blue.

### Subcellular localisation of new regulators of glycoprotein secretion

Given the poor characterisation of GPR161, TMEM220, FAM98B, FAM102B and MXRA7 in the literature and the limited resources available to study their localisation, we obtained cDNA constructs of each of these gene fused to a FLAG-tag, to study the localisation of the protein inside the cells (figure 4). HeLa cells were transiently transfected with cDNA, then fixed 24 h after transfection. Fixed cells were immunostained for FLAG and either calnexin or TGN46, markers of the ER and the Golgi, respectively. GPR161 localised to the Golgi, as seen by its co-localisation with TGN46 (figure 4A), whereas TMEM220 and MXRA7 both partially colocalised with calnexin, indicating that they are found at the ER (figure 4B-C). FAM98B was primarily found in the cytosol, with a portion of FAM98B also localising to the nucleus (figure 4D). Finally, FAM102B was found both at the plasma membrane and in the cytosol (figure 4E). However, it is important to note that the overexpression of these FLAG-tagged construct may not reflect the endogenous protein localisation, as some dominant negative effect might happen when using the cDNA construct. We believed this was the case for TMEM220, as the TGN46 immunostaining seems compromised in the transfected cells (figure 4B). It was therefore important to validate the localisation of the protein using specific antibodies. We studied the localisation of the endogenous proteins GPR161 and TMEM220 and confirmed that GPR161 localised mainly in the Golgi (figure 5Ai). This was further demonstrated using siRNA against GPR161; here the Golgi localisation of GPR161 disappeared (figure 5Bi). However, endogenous TMEM220 localised to the Golgi (figure 5Aii), in contrast to the cDNA overexpression showing an ER localisation. This Golgi residence was similarly confirmed using siRNA (figure 5Bii). It is therefore likely that the overexpression of TMEM220 causes a perturbation in the trafficking of proteins from the ER to the Golgi.

**Figure 4:**
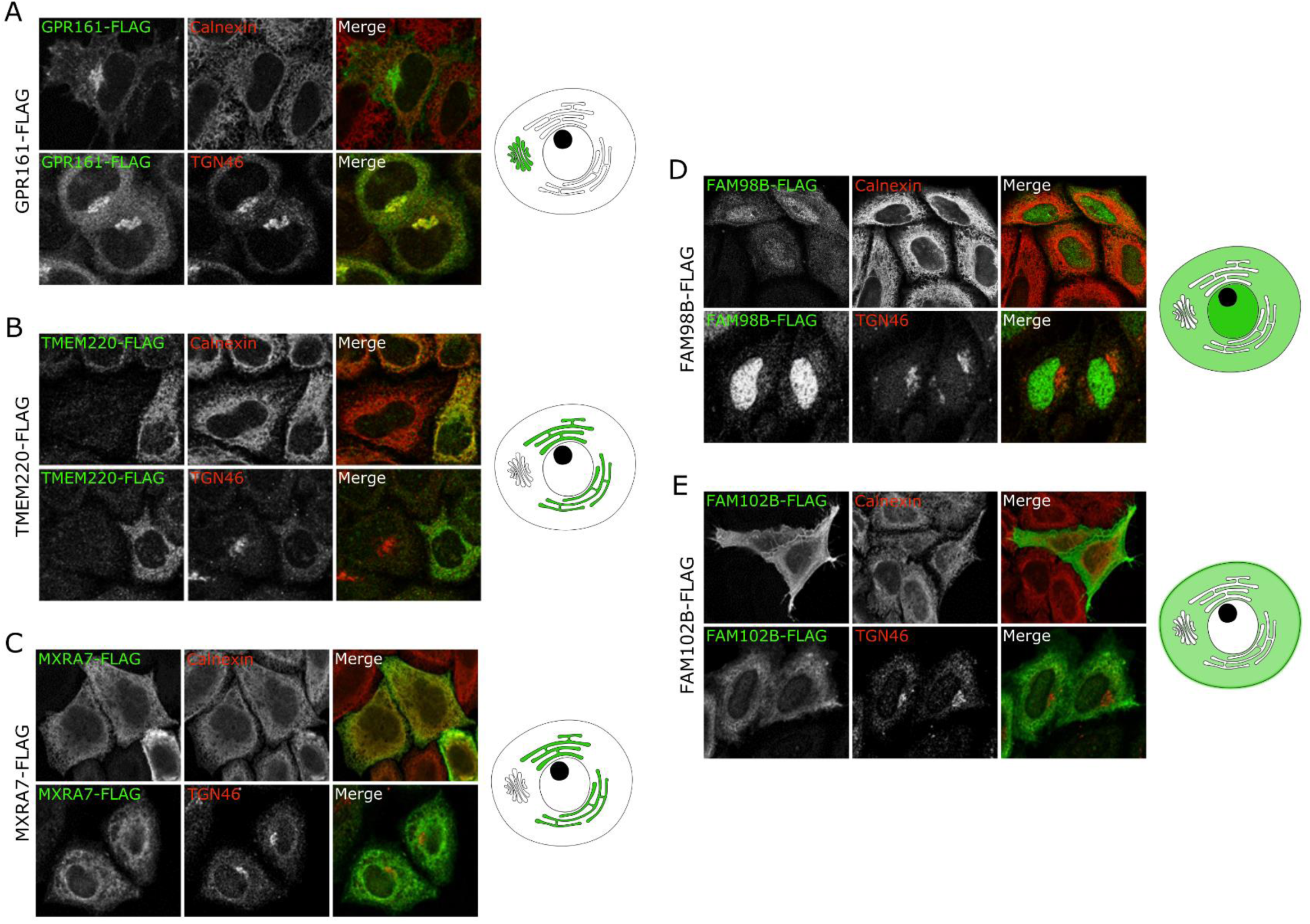
Localisation study for new regulators of glycoprotein secretion. Fluorescence microscopy images of HeLa cells expressing a FLAG-tagged cDNA construct show each protein’s localisation by co-staining with either an ER marker, calnexin, or a Golgi marker, TGN46. In the merged images, FLAG staining is shown in green and calnexin or TGN46 staining is shown in red. The subcellular localisation of each protein construct is indicated in green on the cell diagrams. Darker shades of green are used to indicate the most prominent location if the construct localises to more than one compartment. A-E show expression of FLAG-tagged GPR161, TMEM220, MXRA7, FAM98B and FAM102B.

**Figure 5:**
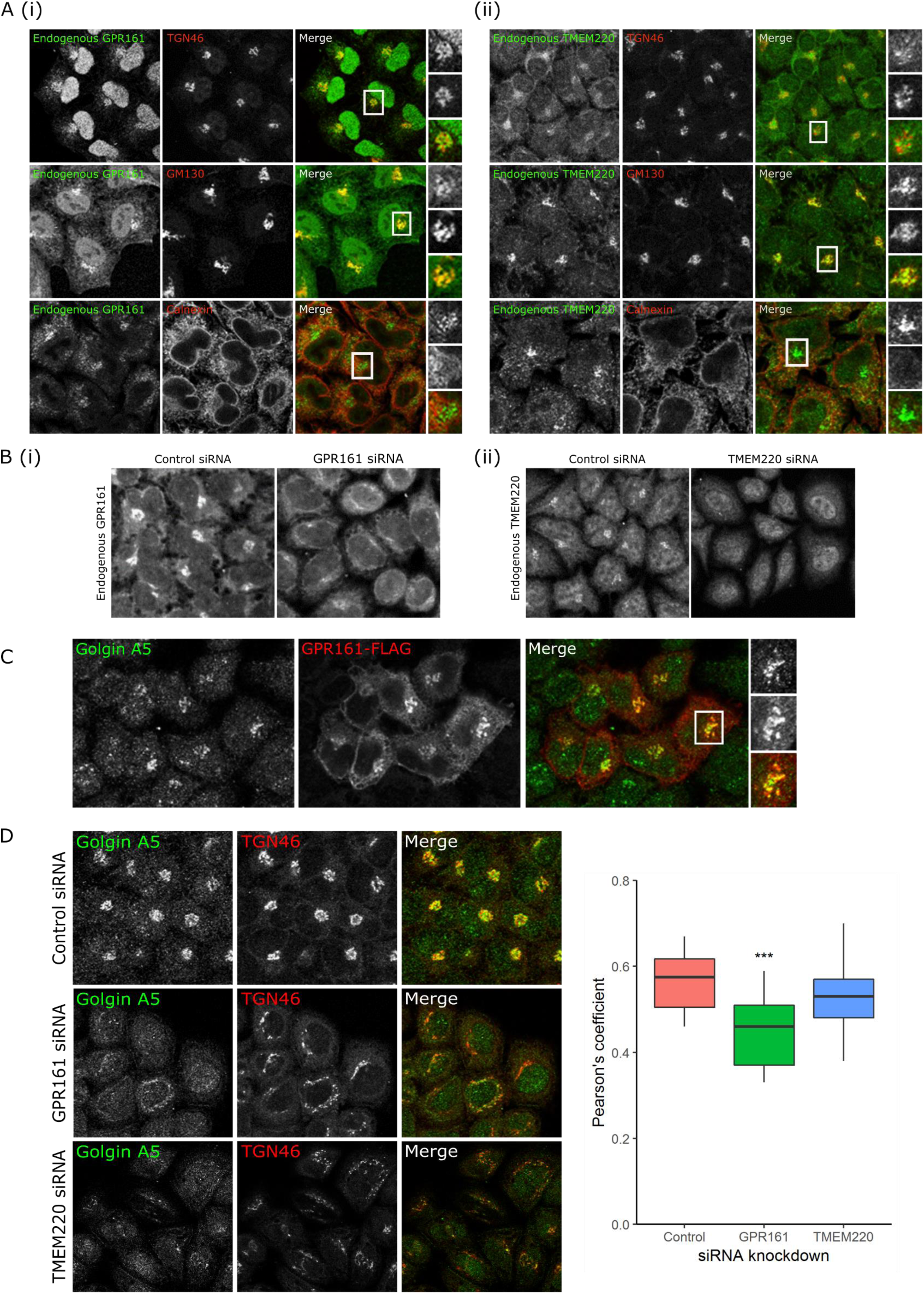
GPR161 and TMEM220, two novel Golgi-resident proteins involved in protein secretion, regulate glycoprotein secretion by different mechanisms. (A) Colocalisation microscopy shows that both (i) GPR161 and (ii) TMEM220 colocalise with TGN46 and GM130, two Golgi markers, and not with calnexin, a marker of the ER. (B) siRNA knockdown of (i) GPR161 or (ii) TMEM220 demonstrates that the antibodies used to detect these endogenous proteins only specifically detect protein at the Golgi. (C) GPR161-FLAG (red) colocalises with golgin A5 by microscopy. (D) Microscopy shows that knockdown of both GPR161 and TMEM220 lead to Golgi fragmentation seen by the TGN46 staining, but only GPR161 knockdown results in less colocalisation of golgin A5 and TGN46. Quantification of this change in colocalisation, measured by Pearson’s coefficient, shows that GPR161 knockdown cells have significantly less colocalisation than control cells (p < 0.001), but there is no significant difference in colocalisation between control cells and TMEM220 knockdown cells (p = 0.302).

### GPR161 is a novel Golgi resident protein involved in protein secretion

Our study shows that GPR161 is a new Golgi-resident protein. To understand its role in secretion, we used the BioPlex network, a resource of proteins shown to interact by affinity purification-mass spectrometry, to look at its interacting partners^20,21^. Interestingly, the BioPlex network shows that GPR161 interacts with golgin A5 (also known as golgin 84). Golgin A5 is a coiled-coil membrane protein that plays a role in intra-Golgi vesicle capture^22^ and contributes to maintaining Golgi morphology, likely by its interaction with Rab1A^23–25^. We showed that golgin A5 and GPR161 colocalised (figure 5C), and the knockdown of GPR161 decreased the colocalisation between golgin A5 and TGN46 (figure 5D). This appears to be specific to GPR161, since the knockdown of TMEM220, which we also found localises to the Golgi, did not affect golgin A5-TGN46 colocalisation, showing that Golgi fragmentation alone does not change golgin A5 localisation to the Golgi (figure 5D). These data suggest that GPR161 may serve to recruit golgin A5 to the Golgi membrane.

## DISCUSSION

Here we identified novel genes involved in glycoprotein secretion, using a combination of pooled CRISPR screening and siRNA screening. This work expands on our results from a previous CRISPR screen measuring cell-surface galectin-3, which had identified a number of regulators of glycosylation and protein trafficking; however, most of those hits had had well-known functions^14^. Although galectin-3 itself is unconventionally secreted, this type of screen should primarily identify genes responsible for the transport of glycoproteins to the cell surface, as secreted galectin-3 can be transferred between cells^14,26^. Here we sought to build on the findings of our previous screen by using galectin-3 to identify novel proteins with roles in glycoprotein secretion. We used the improved Brunello sgRNA library^18^ rather than GeCKOv2 library, and an altered cell sorting strategy which would allow more hits through cell sorting initially, in order to increase the number of true positives in later populations. The Brunello library is an optimised library for the human genome, which was designed to increase the percentage of active guides while reducing off target effects compared to other libraries^18^. Furthermore, here we analysed populations at all stages of cell sorting rounds performed rather than only the final sorted population.

Overall, our data suggest that the enhanced cell sorting strategy we employed here does improve results. While there is little enrichment visible by anti-galectin-3 staining in the earlier sorted populations, deep sequencing showed that enrichment had taken place. Many of the hits enriched in the first sorted population, N1, localise to the ER and the plasma membrane; later populations had fewer hits localised to the ER. This is likely due to problems with cell survival; over the extended time period and stress derived from the four sorts, cells with ER defects may be less able to survive. Enrichment of two distinct galectin-3 negative populations was visible by flow cytometry after the third round of cell sorting, in population N3. The most negative population within N3 consisted of cells that had little or no galectin-3 bound to the cell surface but were still able to secrete galectin-3 into the supernatant, so the excess of extracellular galectin-3 could bind to both the mid-negative population and the unaffected population of cells. Combined with the higher effective concentration of anti-galectin-3 antibody during staining, this meant that the unaffected population appeared to have more galectin-3 staining after earlier sorts. Across all sorted populations, the enrichment of many hits that are known to be in secretory and glycosylation pathways provides an initial validation of the success of the CRISPR screen. In comparison to our previous screen, we identified more hits known to have roles in the maturation or secretion of glycoproteins; moreover, we found a much higher number of genes not known to have such roles.

Due to the large number of hits identified by the MaGECK algorithms, we selected hits to be taken forward for secondary screening for protein secretion. Via this screening method, we identified 92 hits that result in a secretion defect, of which 55 are not annotated with GO terms related to secretion. The 92 hits validated for secretion represent approximately one quarter of the hits screened, highlighting the importance of secondary screening after a CRISPR screen due to the high chance of false positives. However, the fact that 55 of the validated hits have previously unknown or unclear functions in secretion demonstrates the power of this screening strategy.

To further validate these hits, we investigated Golgi morphology. There was a clear increase in the percentage of cells with Golgi fragmentation for many of the hits. As siRNA knockdown efficiency was not quantified for each gene, the percentage of cells with fragmented Golgi does not give a definitive ranking of the scale of the effect each gene has on protein secretion. Additionally, while the presence of fragmented Golgi gives a reason for the altered secretion and therefore validates both the CRISPR screen and HRP assay results, it does not explain how each individual gene product contributes to protein secretion. As such, we selected five hits with little known about their function in protein secretion to investigate further. Most of these had very strong phenotypes in both assays, such as MXRA7, but some had a weaker, yet still significant, phenotype, including GPR161; we studied this selection in order to validate the full list of genes as a resource for further investigation.

Among the five hits that we studied for further mechanistic insight, GPR161 and TMEM220 localised to the Golgi whereas MXRA7 localised to the ER. FAM98B and FAM102B localised to the cytosol or the plasma membrane. Further studies will be required to fully understand how each gene regulates protein secretion. Interestingly, a recent study has shown the importance of the extracellular matrix (ECM) in regulating Golgi organisation and function via the activation of ARF1^27^. MXRA7, a component of the ECM, might play a similar role. Little is known about FAM98B and FAM102B, but FAM102B shows a clear localisation at the plasma membrane, so it could regulate protein secretion through an identified signalling pathway. FAM98B shows cytosolic and nuclear localisation, and interacts with other such proteins to form a complex involved in shuttling RNA between the nucleus and cytoplasm^28^, suggesting that FAM98B may affect protein secretion by a transcriptional route.

Our further investigation of the two proteins that localised to the Golgi, TMEM220 and GPR161, suggests that they regulate Golgi morphology by different mechanisms. TMEM220 is a transmembrane protein that has been shown to interact with both actin and testis-specific glyceraldehyde-3-phosphate dehydrogenase (GAPDH) in the BioPlex network^20,21^. Actin, and the cytoskeleton in general, is well known to contribute to protein trafficking along the secretory pathway; for example, it is recruited to the Golgi to by a complex of SPCA1 and active cofilin, where it is suggested to form a membrane domain required to initiate sorting of secretory cargo^7,19,29^. It is possible that TMEM220 acts as an anchor for actin on the Golgi membrane. This possibility needs to be further investigated using *in vitro* experiments. Alternatively, TMEM220 could affect protein secretion via an interaction with GAPDH, which has recently been implicated in protein secretion via the inhibition of COPI vesicle biogenesis^30^.

GPR161 is a G-protein coupled receptor (GPCR) that is involved in neural tube development and acts as a regulator of cell signalling pathways, including Shh signalling, PKA signalling, retinoic acid signalling and Wnt signalling^31,32^. This was a particularly interesting hit as it has recently become clear that GPCRs function at membranes other than the plasma membrane^33^, and previous work has suggested that a Golgi-resident GPCR regulates transport from the Golgi, although the specific GPCR involved remains unknown^34^. Our results showing that GPR161 is both localised to the Golgi and involved in regulation of Golgi morphology and glycoprotein traffic fit with this emerging idea that GPCRs can regulate protein trafficking. The BioPlex network for GPR161 shows an interaction with golgin A5, as well as with five PKA subunits^20,21^; other work has also demonstrated an interaction between GPR161 and PKA^35^. The interaction with PKA subunits is interesting, as a PKA signalling pathway has previously been shown to regulate retrograde transport from the Golgi to the ER, which indirectly also affects anterograde traffic^36^. However, previous work found that PKA regulatory subunits binding to GPR161 results in its transport to the plasma membrane to signal through PKA^35^. Here, we did not observe any localisation of either overexpressed or endogenous GPR161 at the plasma membrane, suggesting that a different mechanism may be important here. Furthermore, our data suggest that the interaction between GPR161 and golgin A5 is important for maintenance of the Golgi architecture and function, and we propose that GPR161 may act to recruit golgin A5 to the Golgi. To confirm this, further experiments involving mutated forms of GPR161 and golgin A5 will have to be performed.

In summary, here we describe an optimised CRISPR screening strategy that successfully identifies new regulators of glycoprotein secretion. Our secondary screening validated 55 hits not previously known to be directly involved in protein secretion; many of these hits also regulate Golgi morphology. We hope that these validated hits can serve as a resource for other researchers investigating protein secretion and Golgi morphology. We also highlight GPR161, a particularly interesting protein given that Golgi-localised GPCRs have recently been implicated in protein trafficking. We find that it is a novel Golgi-localised protein that appears to interact with golgin A5 in order to maintain Golgi structure.

## MATERIALS AND METHODS

### List of reagents

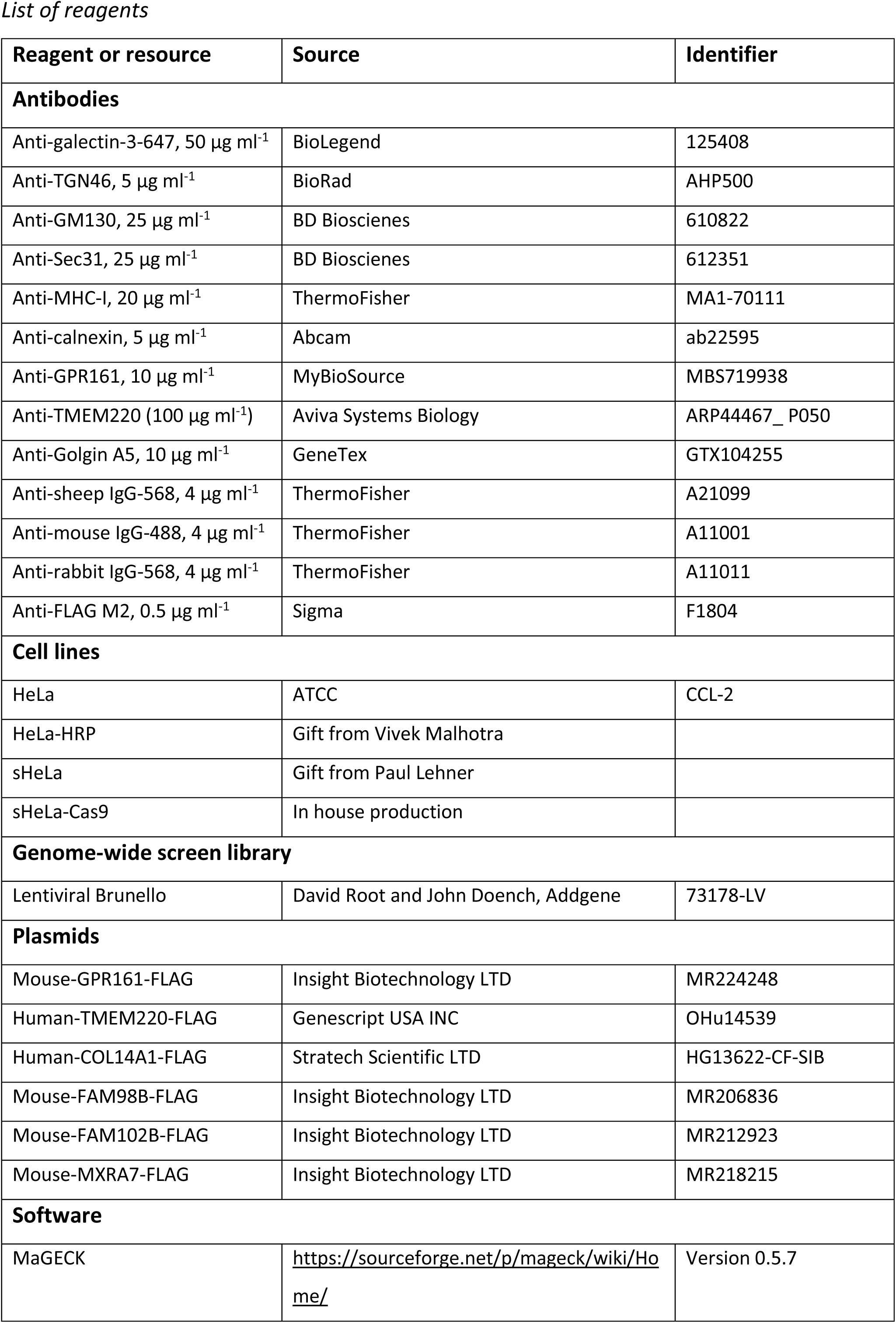

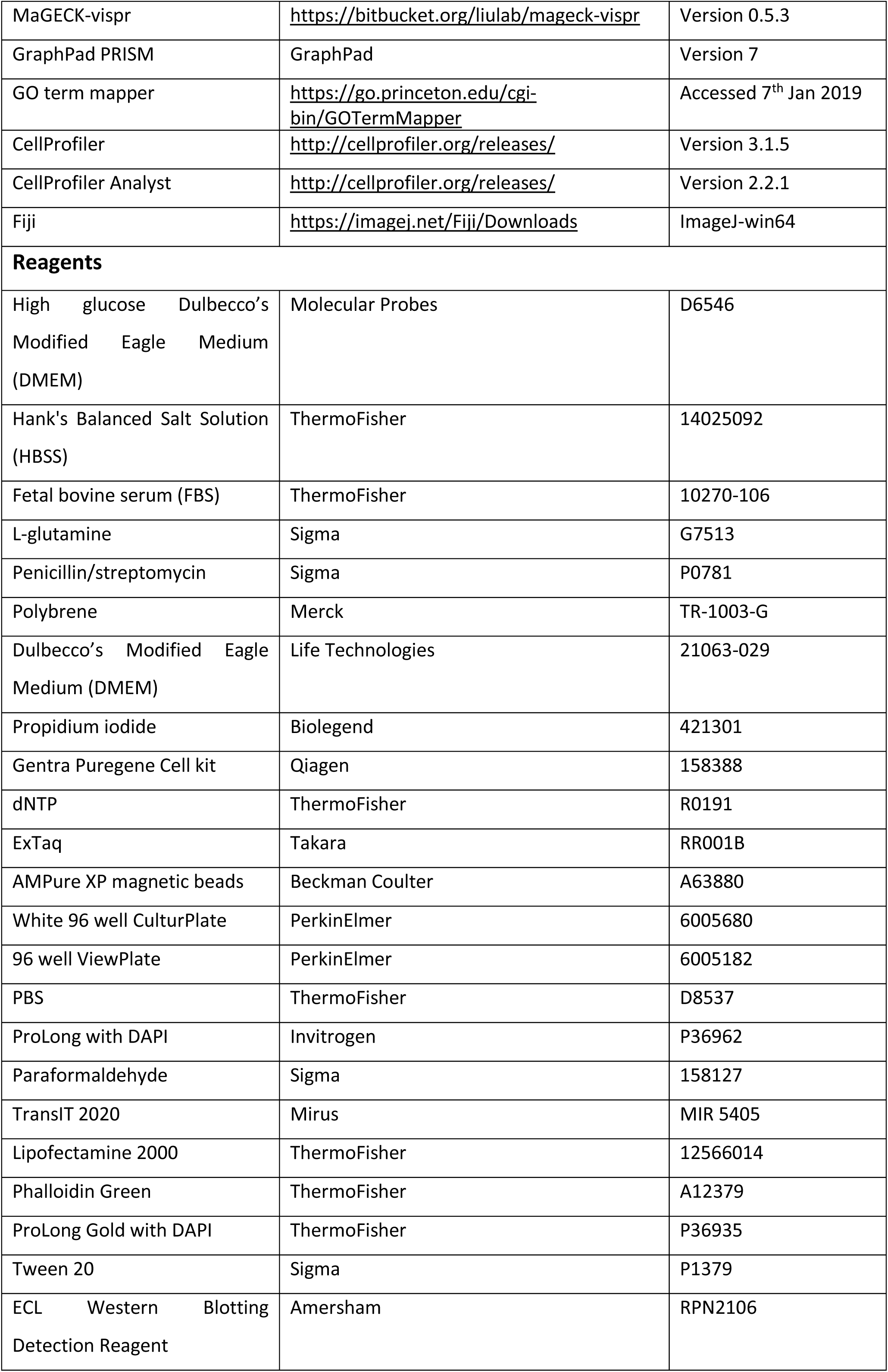

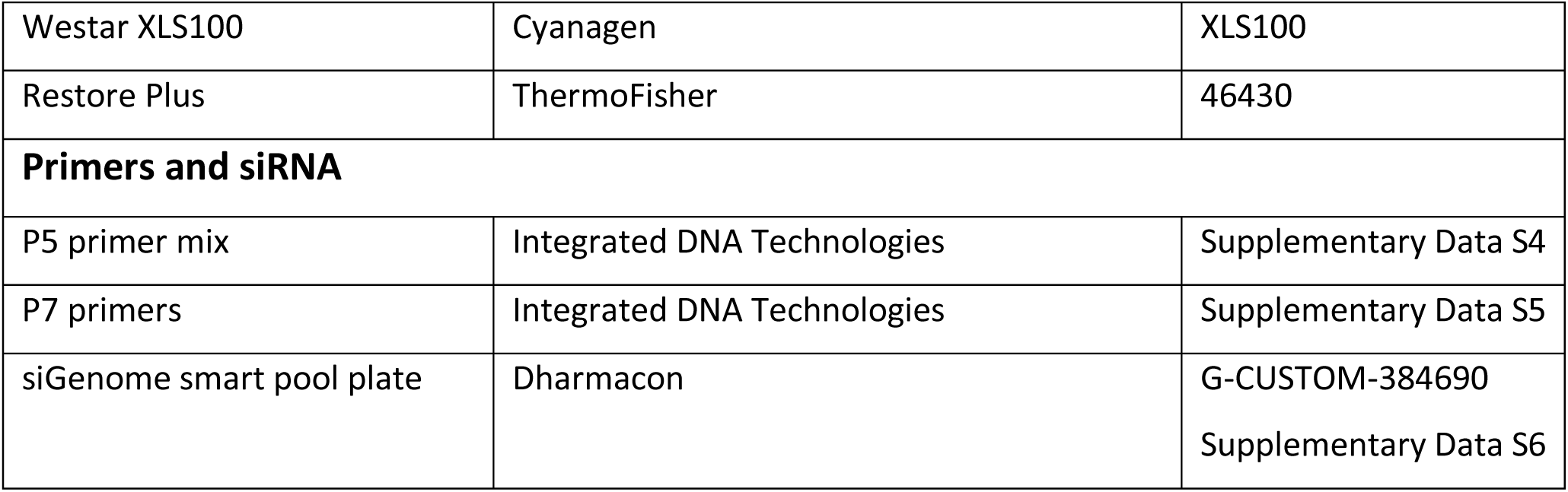

### Cell culture

HeLa cells, suspension HeLa cells expressing Cas9 (sHeLa-Cas9) and HeLa cells expressing horseradish peroxidase fused to a signal sequence to direct its secretion (HeLa-ss-HRP) were cultured in 10% culture medium: 10% (*v/v*) FBS, 100 U Penicillin / 0.1 mg ml^-1^ streptomycin, 2 mM L-glutamine in high glucose DMEM, in 5% CO2 at 37°C.

### CRIPSR screen: transduction with Brunello library

24 h before transduction, sHeLa cells were passaged. The lentiviral Brunello library, which targets 19,114 genes in the human genome with four unique guides per gene,^18^ was used to transduce 1 x 10^8^ suspension HeLa cells expressing Cas9 (sHeLa-Cas9) in polybrene media (10 µg ml^-1^ polybrene, 10% (*v/v*) FBS, 100 U Penicillin / 0.1 mg ml^-1^ streptomycin, 2 mM L-glutamine in high glucose DMEM (Sigma, D6546)) at an MOI of 0.50 by spinoculation at 1000 xg for 30 min at 20°C. This gave a transduction efficiency of 40% and therefore an overall percentage of 31% cells having one guide per cell, as calculated using the Poisson distribution for the probability that a cell will be infected with at least one virus, 1-P(0, MOI)^37^. Cells were incubated for 24 h at 37°C, 5% CO2 before passage into puromycin media (1 µg ml^-1^ puromycin, 10% (*v/v*) FBS, 100 U Penicillin / 0.1 mg ml^-1^ streptomycin, 2 mM L-glutamine in high glucose DMEM). Cells were cultured in puromycin media for seven days to retain only transduced cells.

### CRISPR screen: cell sorting by flow cytometry

Cells were washed once with DMEM, incubated with 50 µg ml^-1^ anti-galectin-3 antibody conjugated to Alexa Fluor 647 for 30 min at 4°C, then washed again. Approximately 3 x 10^7^ cells were sorted at the NIHR Cambridge BRC cell phenotyping hub on a BD FACS AriaIII and FACS Aria Fusion (using a 100um Nozzle and run at a pressure of 25psi); Alexa Fluor 647 fluorescence was detected with a 670/30 BP detector on the AriaIII and Aria Fusion cell sorters. Target cell population to be sorted was gated based on the lowest anti-galectin-3-AF647 fluorescent signal, using Purity as sort precision mode. After cell sorting, cells were cultured in 20% serum media for 24-48 h, then cultured in culture media. When the population had expanded to 5 x 10^7^ cells, two samples were taken; 5 x 10^6^ cells was frozen for guide sequencing, and 1 x 10^6^ was assessed by flow cytometry. Gating leniency was decreased with progressive rounds, as shown in Figure 1. After the third sort, two distinct negative populations were observed so a fourth sort was performed to separate these two populations, gating each population around a 5-10% peak. For flow cytometry of 1 x 10^6^ cells after cell sorting, cells were immunostained as above, with an additional propidium iodide stain for 5 min, and analysed on an Accuri™ C6 (BD Biosciences) equipped with lasers providing 488 nm and 640 nm excitation sources. Alexa Fluor 647 fluorescence was detected with an FL4 detector (675/25 BP).

### CRISPR screen: Library preparation and sequencing

To prepare samples for sequencing, genomic DNA from the different cell populations was extracted from frozen cell pellets using Gentra Puregene Cell kit and concentration of DNA measured on the nanodrop. PCR was carried out in quadruplicate to amplify an amplicon containing the guide, with primers used to attach barcodes, stagger regions and sequencing adaptors for use in sequencing, as previously described by Doench *et al*^18^. Briefly, each well was set up to contain 10 µg genomic DNA, 0.5 µM uniquely barcoded P7 primer, 0.5 µM P5 stagger primer mix, 200 µM each dNTP, 7.5 units ExTaq and 1x ExTaq buffer in a total volume of 100 µl. PCR cycles were: initial denature step at 95°C for 1 min; 28 cycles of 95°C for 30 s, 53°C for 30 s, 72°C for 30s; final extension at 72°C for 10 min. One replicate from each sample was analysed on a 2% agarose gel to confirm amplification was successful. PCR products were pooled, adding 30 µl from each PCR reaction, then purified using AMPure XP magnetic beads to retain only DNA fragments larger than 100 bp^18^. An equal volume of the pooled product and AMPure XP magnetic beads were mixed at room temperature for 5 min, then placed on a magnet to retain beads. Beads were washed three times with 70% ethanol and purified PCR product was eluted with 500 µl EB buffer. Sequencing was carried out by the CRUK genomics facility using SE50 sequencing on the Illumina HiSeq 4000.

### CRISPR screen: Bioinformatics analysis using MaGECK

Data were analysed using both MaGECK-MLE^38^ and MaGECK-RRA^39^, using an enrichment (positive) sort type. In all analyses, median normalisation was used. PCR replicates were treated as technical replicates and sort replicates were treated as biological replicates. Each population was given an identification code; descriptions are available in supplementary data S2. Genes were annotated with their function using UniProt^40^, and with the organelle they reside in using Gene Ontology subsets^41,42^.

### Secondary screen: HRP assay

Based on having an unknown or unclear function in secretion, hits from the MaGECK analyses were selected to create a shortlist of 368 genes to be used in a secondary screen. This shortlist was screened in an arrayed siRNA screen, using siGenome smart pools (Dharmacon) in which each plate contained two replicates of non-targeting siRNA as a negative control. To screen for conventional secretion, HeLa cells expressing ss-HRP were reverse transfected in with the secondary screen library as previously described^43^. Briefly, 80 µl of 31 000 cells ml^-1^ were plated into wells of a 96 well plate containing a mixture of 20 µl 250 nM siRNA and 0.5% (*v/v*) lipofectamine 2000 in OptiMEM and incubated at 37°C, 5% CO2. Medium was changed to fresh culture medium 48 h after transfection. 72 h after transfection both supernatants and cell lysates were assessed for HRP levels in white CulturPlate 96 well plates by chemiluminescence using a Tecan M1000 Pro. Luminescence signal from the cell lysate (CL), supernatant (SN) and the CL:SN ratio were each normalised to signal obtained from siRNA control cells according to the formula:

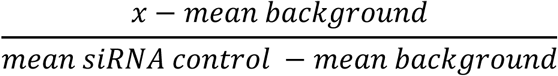

To identify genes required for HRP secretion, a line with a gradient of 1 was fit to the normalised CL vs normalised SN data. Outliers of this line, as identified by the ROUT method^44^, are genes that result in an up or down regulation of HRP secretion. The same method was also used to identify outliers from the line of y = 1 for the normalised CL:SN ratio line. The generic gene ontology (GO) term mapper was used to classify hits identified into broad categories^45^.

### Secondary screen: Golgi morphology

HeLa cells were reverse transfected as described above, although here transfected in 96 well ViewPlates. 72 h after transfection cells were fixed in ice cold methanol for 5 min, washed in PBS, stained with appropriate primary and secondary antibodies and ProLong with DAPI, and observed using an Opera Phenix High-Content Screening System to obtain unbiased confocal pictures. Analysis was performed on CellProfiler to define cells and Golgi, and to measure Golgi intensity, size, shape and granularity. The classifier of CellProfiler Analyst (CPA) was then trained using cells randomly chosen from the whole experiment. These were classified as having fragmented or intact Golgi until the sensitivity reached an acceptable level: 82% and 77% for the intact Golgi and fragmented Golgi classes respectively; here 65 cells were used. The CPA classifier was then used to define all cells as having intact or fragmented Golgi.

### Transfection with cDNA

HeLa cells at ∼50% confluency were transfected with a mix of cDNA at 1 µg µl^-1^ and 0.2% (*v/v*) TransIT in OptiMEM. Cells were incubated at 37°C, 5% CO2 for 4 h, then medium was changed to fresh culture medium; cells were then incubated for a further 20 h.

### Immunofluorescence microscopy

For immunofluorescence microscopy, cells were cultured on coverslips, fixed with either 4% paraformaldehyde in PBS for 5 min and permeabilised with 0.1% Triton X100 in PBS for 5 min, or fixed and permeabilised with ice cold methanol for 5 min. Cells were then blocked in 10% (*v/v*) FBS / 1x PBS for 30 min, incubated with primary antibodies for 2 h, washed three times with PBS, and incubated with secondary antibodies for 30 min. Samples were mounted using ProLong Gold antifade reagent with DAPI and observed using a Leica SP8 laser confocal microscope.

### Colocalisation analysis

To measure colocalisation between golgin A5 and TGN46, 17 cells were selected from each treatment from confocal micrographs and analysed using the coloc 2 plugin in Fiji^46^. For cells with TMEM220 or GPR161 knocked down, only cells with fragmented Golgi were selected from confocal micrographs to be confident that siRNA knockdown was effective. Pearson’s coefficient with no threshold was recorded for each cell analysed. Data were tested for normality and equality of variance by the Shapiro-Wilk test and Bartlett’s K squared test respectively. After passing normality tests, data were analysed by one-way ANOVA followed by Tukey HSD post-hoc test between groups.

## ACKNOWLEDGMENTS

We thank Paul Lehner for sHeLa cells, Vivek Malhotra for HeLa-ss-HRP cells, David Root and John Doench for the Brunello library, Marcella Ma and Giles Yeo for valuable help and advice with the gRNA sequencing, Matt Castle for advice on statistical analysis, and James Smith and Greg Strachan for technical assistance with the Opera Phenix.

## COMPETING INTERESTS

The authors declare no competing or financial interests.

## AUTHOR CONTRIBUTIONS

Conceptualisation: S.J.P., J.V. and K.M.; Experimental work: S.J.P., J.V., E.P.G. and K.M.; Validation: S.J.P., J.V. and K.M.; Formal analysis: S.J.P. and K.M.; Writing – original draft: S.J.P., S.E.S., J.V. and K.M.; Supervision: K.M. and S.E.S.; Project administration: K.M.; Flow cytometry administration: A.P.H.; Funding acquisition: S.J.P., A.P.H., S.E.S. and K.M.

## FUNDING

This work was supported by a Wellcome Trust Strategic Award [100574/Z/12/Z], a Wellcome Trust PhD studentship for S.J.P., the Medical Research Council Metabolic Diseases Unit [MRC_MC_UU_12012/5], the Isaac Newton Trust/Wellcome Trust ISSF/ University of Cambridge joint research grant for J.V. and K.M., Biotechnology and Biological Science Research Council Future Leader Fellowship for S.E.S and the National Institute for Health Research for E.P.G. and A.P.H.

