## Supplementary Info 3 for "Genome-wide CRISPR screening identifies new regulators of glycoprotein secretion"

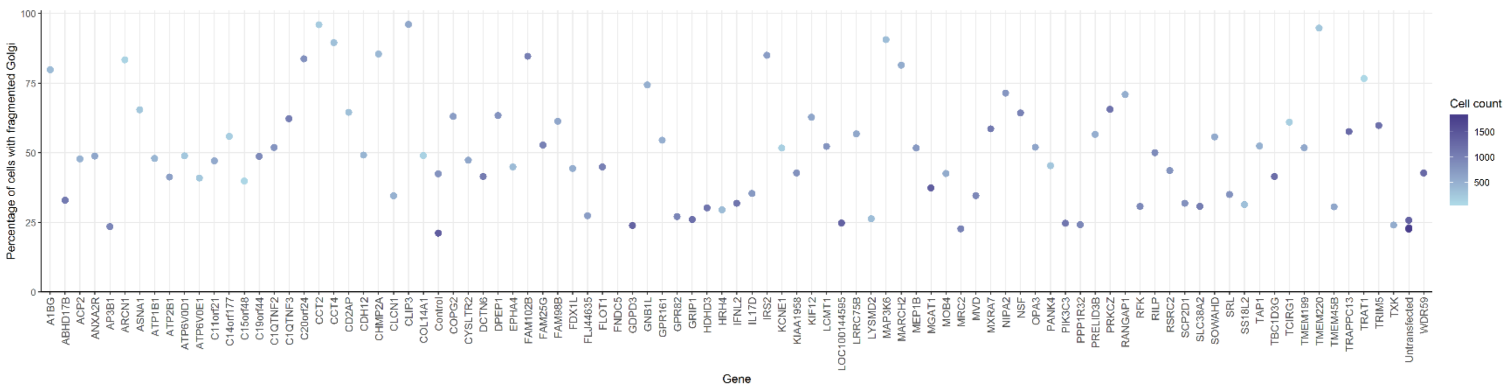


**Supplementary Figure S3: All hits from the secondary screen for Golgi morphology.** Percentage of cells with fragmented Golgi for all of the hits screened in the tertiary screen. As in figure 3, hits are arranged alphabetically and coloured by cell count, with darker blue spots representing more confluent wells.
